# Comparative genomic analysis of skin and soft tissue *Streptococcus pyogenes* isolates from low- and high-income settings

**DOI:** 10.1101/2021.09.10.459590

**Authors:** Saikou Y. Bah, Alexander J. Keeley, Edwin P. Armitage, Henna Khalid, Roy R. Chaudhuri, Elina Senghore, Jarra Manneh, Lisa Tilley, Michael Marks, Saffiatou Darboe, Abdul K. Sesay, Thushan I de Silva, Claire E. Turner, on behalf of The MRCG StrepA Study Group

## Abstract

*Streptococcus pyogenes* is a leading cause of human morbidity and mortality, especially in resource limited settings. The World Health Organisation has recently made a vaccine for *S. pyogenes* a global health priority to reduce the burden of the post-infection rheumatic heart disease. For a vaccine to be active against all relevant strains in each region, molecular characterisation of circulating *S. pyogenes* isolates is needed. We performed extensive comparative whole genome analyses of *S. pyogenes* isolates from skin and soft tissue infections in The Gambia, West Africa, where there is a high burden of such infections. To act as a comparator to this low-income country (LIC) collection of isolates, we performed genome sequencing of isolates from skin infections in Sheffield, UK, as representative high-income country (HIC) isolates. LIC isolates from The Gambia were genetically more diverse (46 *emm*-types in 107 isolates) compared to HIC isolates from Sheffield (23 *emm*-types in 142 isolates), with only 7 overlapping *emm*-types and with diverse genetic backgrounds. Characterisation of other molecular markers indicated some shared features, including a high prevalence of the skin infection-associated *emm*-pattern D and the variable fibronectin-collagen-T antigen (FCT) types FCT-3 and FCT-4. A previously unidentified FCT (FCT-10) was identified in the LIC isolates, belonging to two different *emm*-types. A high proportion (79/107; 73.8%) of LIC isolates carried genes for tetracycline resistance, compared to 53/142 (37.3%) HIC isolates. There was also evidence of different circulating prophages, as very few prophage-associated DNases and lower numbers of superantigens were detected in LIC isolates. Our study provides much needed insight into the genetics of circulating isolates in a LIC (The Gambia), and how they differ from those circulating in HICs (Sheffield, UK). Common molecular features may act as bacterial drivers for specific infection types, regardless of the diverse genetic background.

## Introduction

*Streptococcus pyogenes* (Group A *Streptococcus*, GAS) is a human-specific pathogen and a leading cause of morbidity and mortality, especially in resource-limited countries. *S. pyogenes* can cause diseases ranging from mild superficial infections, such as impetigo and pharyngitis, to invasive diseases such necrotising fasciitis and streptococcal toxic shock syndrome (1) and can also cause post-infection autoimmune sequalae such as acute rheumatic fever (ARF) leading to rheumatic heart disease (RHD). There is a substantial global burden of RHD, accounting for approximately 320,000 deaths in 2015, the majority of which were recorded in sub-Saharan Africa (2). Recognising this burden, the World Health Organisation (WHO) has prioritised the need for a vaccine that would have global coverage, and recommended an increase in research, especially in low- to middle-income countries (LMICs) (3).

Progress towards a vaccine for *S. pyogenes* has been hampered over the years by the association of the most promising vaccine candidate, the surface protein M, with the development of RHD. This may be circumvented by targeting the N terminal portion of the M protein, but this region is hypervariable, thus any vaccine would be serotype/genotype specific. *S. pyogenes* isolates are genotyped by sequencing the corresponding hypervariable 5’ region of the M protein-encoding gene, *emm*. Over 220 different *emm*-types have been identified globally, but in high income countries (HICs) the majority of disease is caused by a limited number of *emm*-types, with *emm*1 being the most common. A 30-valent M-protein vaccine has been developed and is undergoing clinical trials but is based on genotypes predominantly circulating in Europe and North America (4, 5). The limited available data for *S. pyogenes* in LMICs suggest a far more genetically diverse population than that seen in HICs (6–9). More extensive, global genomic analysis may reveal another vaccine target or combination of targets that would be applicable in these settings.

The *emm* gene lies within the *S. pyogenes* core *mga* (multi gene activator) regulon locus, and upstream and/or downstream of the *emm* gene there may be additional *emm*-like genes. There have been ten different *emm* patterns identified, based on the genes within the *mga* regulon, which form three main groupings: A-C, D or E. The majority of *emm* types have been associated with only one *emm* pattern (10). There is some epidemiological evidence supporting the existence of tissue tropism among *emm* types, with preference for either pharyngeal or skin infection sites, or “generalists” that are equally able to infect both sites (11). There is also an association with this tissue tropism to *emm* pattern; pharyngeal specialists are pattern A-C, the skin specialists are pattern D, and the generalists are pattern E (10). However, much of this evidence comes from population-based surveys where there is greater sampling of pharyngeal infections in HICs but more skin infections (impetigo/pyoderma) in LMICs (12). Whether this reflects an actual difference in the prevalence of infection types is unclear, as data is lacking for both skin infections in HICs and pharyngeal infections in LMICs (12–14).

It is estimated that more than 162 million children have impetigo/pyoderma at any given time, predominantly in LMICs, although data for Europe, South-East Asia and North America is very limited (14). A recent study in The Gambia, West Africa, identified a 17.4% prevalence of pyoderma in children, with *S. pyogenes* as a leading infection cause (15). This was higher than the estimated global prevalence of 12.3% (14). The association with scabies infestation in The Gambia was lower than that seen in other settings, but there was an increase in pyoderma prevalence from 8.9% to 23.1% during the rainy season (15). Whilst there may be environmental and socio-demographic factors underpinning the high burden of pyoderma in The Gambia, there may also be bacterial factors involved and potentially tissue tropism. To investigate this, and to provide molecular characterisation of *S. pyogenes* causing skin infections in The Gambia, we performed whole genome sequencing on the isolates obtained from our previous study (15). To act as a comparative HIC collection of isolates, we also performed whole genome sequencing and molecular characterisation of *S. pyogenes* isolated from skin infections in Sheffield, UK. Our study highlights the genetic diversity observed in an LMIC *S. pyogenes* population compared to a HIC population with limited overlap of *emm*-types. However, there were some shared molecular markers associated with skin infection isolates, including *emm*-pattern, *emm*-cluster and FCT-type, supporting the hypothesis that there are bacterial factors driving certain types of infection.

## Material and Methods

### Isolates

*S. pyogenes* skin pyoderma lesion isolates from one hundred and thirty-six children under the age of 5 in the peri-urban setting of Sukuta in The Gambia, collected between May and September 2018 (15), were available for whole genome sequencing. As previously described, swabs were stored in liquid Amies transport medium before being taken to Medical Research Council Unit The Gambia at London School of Hygiene & Tropical Medicine (MRCG at LSHTM) for culture and identification of *S. pyogenes* (15). To provide a representative collection of *S. pyogenes* from a HIC for comparison, 160 sequentially cultured skin and soft tissue infection (SSTI) isolates were collected from the Department of Laboratory Medicine, Northern General Hospital, Sheffield, UK between January and April 2019. No patient data were obtained for these isolates so no selection was applied for patient characteristics such as age or sex.

### Whole genome sequencing

Streptococcal DNA was extracted from isolates using a method previously described (16). For Gambian isolate DNA, sequencing libraries were prepared using the NEBNext Ultra™ II DNA Library Prep Kit for Illumina and sequenced on an Illumina MiSeq at MRCG. The MiSeq V3 reagent kit was used to generate 250bp paired end reads following the Illumina recommended denaturation and loading recommendations which included a 5% PhiX spike-in. Raw sequence quality assessment was performed using FastQC (v0.11.8; https://www.bioinformatics.babraham.ac.uk/projects/fastqc) with default settings and reads were trimmed using Trimmomatic (v0.38) with following settings: LEADING:3 TRAILING:3 SLIDINGWINDOW:4:15 MINLEN:36 (17). Sequencing of the genomic DNA from Sheffield, UK collection isolates and a selection of isolates from the Gambia collection that were subjected to repeat sequencing after failing quality control, was provided by MicrobesNG (microbesng.com) using the Nextera XT Library Prep kit (Illumina) and the Illumina HiSeq/NovaSeq platform generating 250bp pair end reads. Data was subjected to MicrobesNG quality control and Trimmomatic pipelines. Short read sequence data were submitted to the sequence read archive and accession numbers are provided in Supplementary Table 1.

### Whole genome sequence analysis

*De novo* assembly was performed using SPAdes (v3.13.1) with k-mers sizes of 21, 33, 55 and 77 (18). Assembly qualities statistics were generated using Quast (19) (Supplementary Table 1) and any assemblies with more than 500 contigs and a total genome size greater than 2.2Mb were removed from downstream analyses. Prokka (v1.13.3) was then used to annotate the assemblies (20) and the pangenome determined using Roary (v3.12.0) with a 95% identity level (21). Single nucleotide polymorphism (SNP) distances were determined from the Roary core-gene alignment output using snp-dists (v0.7.0, https://github.com/tseemann/snp-dists). RAxML (v8.2.12) (22) was used to generate maximum likelihood phylogenetic trees based on the core-gene alignment, with GTR substitution model and 100 bootstraps. Phylogenetic trees were visualized and annotated using iTOL (23). The *emm* types were determined from the *de novo* assemblies using emm_typer.pl (github.com/BenJamesMetcalf/GAS_Scripts_Reference). Where necessary, *emm* genes were manually located and type determined using the CDC *emm*-typing database (www.cdc.gov/streplab). New *emm*-subtypes were submitted to the database for assignment. Multi-locus sequence types (MLSTs) were determined using the MLST database (pubmlst.org/spyogenes) and a script from the Sanger pathogen genomics group (github.com/sanger-pathogens/mlst_check). Any new alleles and sequence types were submitted to the PubMLST database.

### Variable factor typing

The presence of superantigens; *speA*, *speC*, speG, *speH*, *speI*, *speJ*, *speK*, *speL*, *speM*, *speQ*, *speR*, *ssa* and *smeZ*, and DNases; *sda1*, *sda2*, *sdn*, *spd1*, *spd3* and *spd4* were initially determined by BLAST of representative gene sequences against the *de novo* assemblies. Gene presence was then additionally confirmed by BWA-MEM (24) mapping of the short read sequences to a pseudo sequence of concatenated superantigen and DNase genes; coverage of at least 10 reads across the whole gene was used to confirm presence. Where the BLAST and mapping did not agree, results were manually inspected in the annotated *de novo* assemblies. Antimicrobial resistance (AMR) gene carriage was determined with ABRicate v0.8.13 (github.com/tseemann/abricate) using the ARG-ANNOT database (25), setting a minimum coverage of 70% and percentage identity of 75%.

The nucleotide sequences for *covR*, *covS* and *rocA* regulatory genes and the *hasA*, *hasB* and *hasC* capsule biosynthesis genes from the *S. pyogenes* H293 reference genome (*emm*89, NZ_HG316453.1) were used as queries in blastn searches against the *de novo* assemblies. The start and end coordinates of the best BLAST hits were converted into BED files and used to extract the nucleotide sequences from the *de novo* assemblies using BEDTools (v2.27.1) (26). Extracted gene sequences were then translated into amino acids and variants determined in comparison to the corresponding amino acid sequences of the reference (H293) protein sequences. For *hasABC*, only nonsense variants and gene absence were recorded (Supplementary Table 1).

### *Emm* pattern and FCT regions

To determine the *emm* pattern in the genome of each isolate, *in silico* PCR (https://github.com/simonrharris/in_silico_pcr) was used to extract the sequence of the whole *mga* regulon (the beginning of *mga* to the end of *scpA*) from *de novo* assemblies and then annotated with Prokka. To improve assemblies where the *mga* regulon was not within contiguous sequence, *de novo* assemblies were ordered against a completed reference genome of the same *emm*-type (where available) using ABACAS (27) and the *in silico* PCR repeated. An *emm* pattern of I, II, III, IV, V or VI was assigned using BLAST to identify genes followed by visual determination of gene location within the regulon. For 22 LIC and 3 HIC isolates, the *emm* pattern could not be determined as contiguous sequence for the *mga* regulon could not be obtained (detailed in Supplementary Table 1).

Alleles of the *emm*-like genes *enn* and *mrp* were assigned by comparison to those identified by Frost *et al*. (28), ensuring 100% nucleotide identity across the entire gene sequence. Where we could not obtain contiguous sequence for the *mga* regulon, *enn* and *mrp* alleles were determined by BLAST of each allele sequence against the entire *de novo* assembly. New alleles for *enn* and *mrp* were kindly assigned by Prof Pierre Smeesters and Dr Anne Botteaux. In some cases, breaks in the *de novo* assemblies occurred within the *enn* gene and therefore alleles could not be confirmed (detailed in Supplementary Table 1).

To determine the arrangement of the genes in the FCT region and the FCT type, *in silico* PCR was used to extract the FCT region and annotated with Prokka. Assemblies in which amplicons were not obtained due to contig break in the FCT regions, were again ordered against a close reference of the same *emm*-type (where available). The ORFs within each extracted FCT region were blasted against the entire NCBI database and, in combination with order of the genes, the FCT types were assigned based on previously assigned FCT type where possible (29). For some isolates, it was not possible to obtain contiguous sequence for the FCT region and so the FCT type was estimated based on manual inspection of the *de novo* assembly and identification of FCT associated genes through BLAST.

## Results

### Genetic diversity of *S. pyogenes* LIC and HIC skin and soft tissue isolates

We performed whole genome sequencing on 115 of 127 *S. pyogenes* skin infection isolates collected in The Gambia (15). After quality control and filtering of reads and *de novo* assemblies, we obtained high quality genome sequence data for a total of 107 Gambian (LIC) *S. pyogenes* isolates for further analyses (Supplementary Table 1). Within the genomes of these 107 isolates, we determined 46 different *emm*-types, with no obvious dominant *emm*-type; the most common being *emm*80 (6/107, ~6%), closely followed by *emm*85, *emm*229 and *emm/*stG1750 (5/107 isolates, ~5% each). Although *emm*/stG1750 has been previously identified in group G streptococci, in this case these isolates were *S. pyogenes* with the group A carbohydrate. The multi-locus sequence types (STs) for all 107 isolates were determined and revealed 57 different types, of which 25 were assigned for the first time. Although multiple STs could be found within single *emm*-types, no STs were shared by multiple *emm*-types.

An *emm*-pattern could be assigned to the majority of isolates using the previously determined classifications. The exceptions were two *emm*147 isolates, one *emm*162 isolate, one *emm*247 isolate and five *emm*/stG1750 isolates, for which an *emm* pattern had not been previously described. For the 98 isolates with known *emm*-patterns, 48% (n=47) were D, 40% were E (n=39) and 12% (n=12) were A-C (Figure 1 and Figure 2). In addition to *emm*-pattern, an *emm*-cluster type could also be assigned to these 98 isolates. The *emm*-cluster type is based on the sequence of the full M protein and is broadly associated with *emm*-pattern (30). The majority of isolates (56/98, ~57%,) were assigned to one of the six E *emm*-cluster types: E1 (n=4), E2 (n=2), E3 (n=14), E4 (n=16), E5 (n=2), E6 (n=18), representing 25 *emm*-types. All E1-E4 and all but four E6 *emm*-types were positive for the serum opacity factor (*sof*) gene, commonly associated with E *emm*-clusters (11), however E5 *emm*-types were *sof* negative. The remaining isolates were A-C4 (n=6), D1 (n-1), D2 (n=1), D4 (n=17) or singletons (n=17).

**Figure 1:**
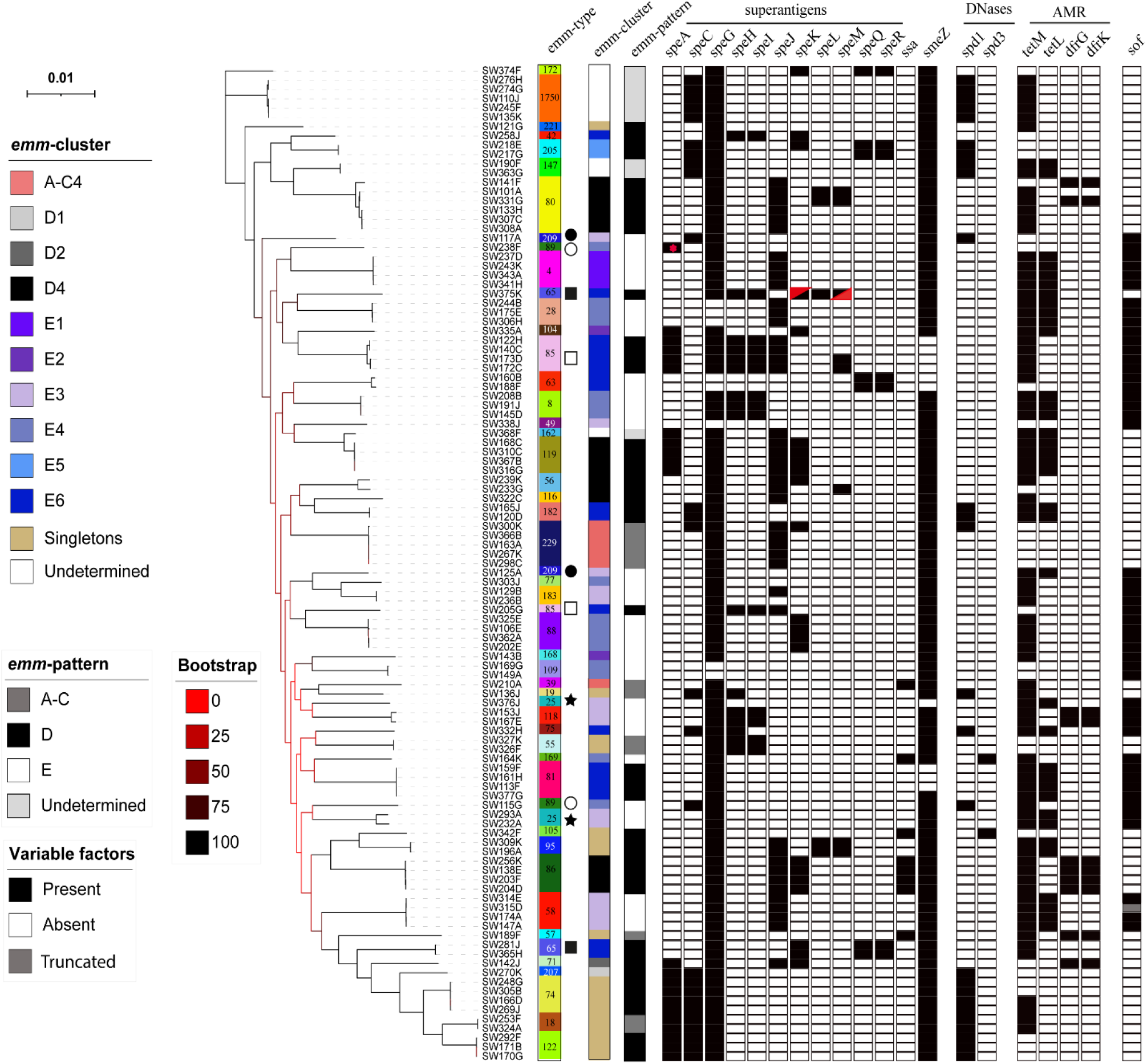
Phylogenetic analysis of 107 genomes from LIC isolates. A maximum likelihood phylogeny was constructed from the core-gene alignment (1,242,112bp) using RAxML (22) with 100 bootstraps. Isolates clustered by *emm*-type except those indicated, whereby two lineages were represented by a single *emm* genotype: star; *emm*25, filled square; *emm*65, open square; *emm*85, open circle; *emm*89, filled circle; *emm*209). Also shown is the presence (black)/absence (white) of the superantigen genes (*speA*, *speC*, *speG*, *speH-M*, *speQ*, *speR*, *ssa* and *smeZ*) and DNase genes *spd1* and *spd3*; four other DNase genes (*sda1*, *sda2*, *sdn*, and *spd4*) were tested for but were not found in any isolate. In all cases, except one (red dot), *speA* was located within the prophage-like element Φ10394.2. One isolate had a gene that appeared to be a fusion of 5’ *speK* and 3’ *speM* (red triangles). Antimicrobial resistance genes (AMR) *tetM, tetL, dfrG* and *dfrK* were also identified in some isolates (white; absent, black; present). The positivity for serum opacity factor (*sof*) is also shown, although for one *emm*55 isolate this gene would produce a truncated variant of SOF (grey). Scale bar represents substitutions per site. *emm*-types are coloured for easy visualisation and type numbers are also given.

**Figure 2.**
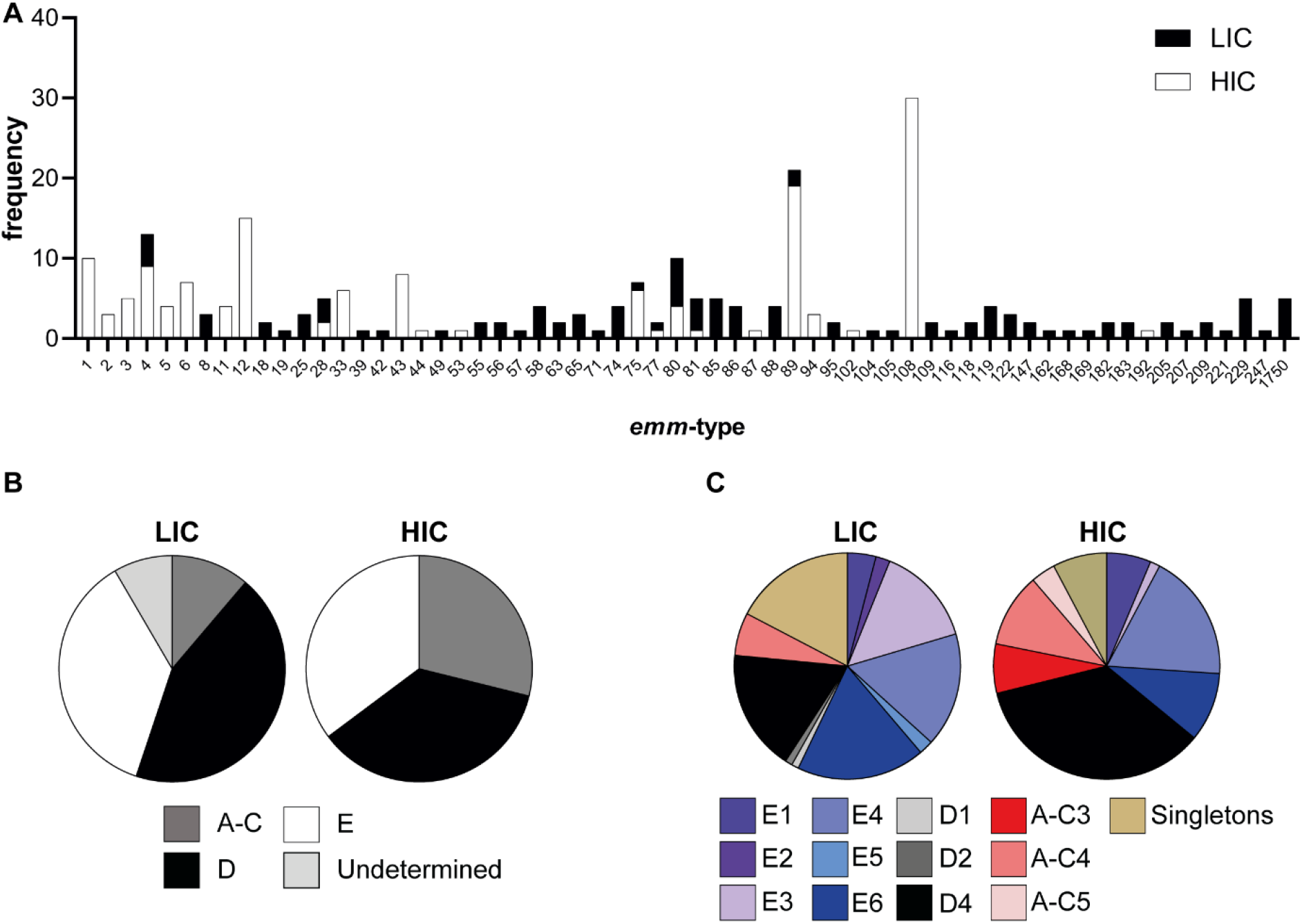
Distribution of *emm*-type, pattern and cluster differs by site. (**A**) The frequency of each of the 62 *emm*-types identified in the LIC isolates (Black) and the HIC isolates (White). (**B**) An *emm*-pattern of A-C, D or E was assigned to 98/107 LIC isolates (the remaining 9 were undetermined) and all 142 HIC isolates. (**C**) An *emm*-cluster was also assigned to 98/107 LIC isolates (the remaining 9 were excluded) and all 142 HIC isolates. Pie charts represent the percentage of isolates associated with each pattern/cluster.

Phylogenetic analysis of the core genome of all 107 LIC isolates showed clustering by *emm*-type (Figure 1). The exceptions to this were *emm*25, *emm*65, *emm*85, *emm*89 and *emm*209, whereby two distinct lineages were identified within these genotypes. Pairwise distance analysis identified a median of 22 SNPs when comparing isolates with the same *emm* type (range; 0-11,142 SNPs), and a median of 9,816 SNPs when comparing isolates with different *emm* types (range 1,423 to 12,428) (Supplementary Figure 1A).

After read quality filtering and assembly assessment, we obtained draft genomes from 142 *S. pyogenes* skin infection isolates collected in Sheffield, UK. Within these 142 HIC isolates there were 23 different *emm*-types but ~59% of the isolates were represented by just 5 *emm*-types: *emm*108 (30/142, 21%), *emm*89 (19/142, 13%), *emm*12 (15/142, 11%), *emm*1 (10/142, 7%) and *emm*4 (9/142, 6%). An *emm*-pattern could be assigned to all 142 isolates and 36% (n=51) were D, 35% (n=50) were E and 29% (n=41) were A-C (Figure 2 and Figure 3). An *emm*-cluster type was also assigned to all 142 isolates and the majority of isolates were D4 (n=50, 35%). No other D cluster-types were found. The most common E cluster type was E4 (n=26), followed by E6 (n=14), E1 (n=9) and E3 (n=2) (Figure 2). The A-C clusters were represented by *emm*1 (A-C3, n=10), *emm*12 (A-C4, n=15) and *emm*3 (A-C5, n=5), which were absent *emm*-types in the LIC population (Figure 2). Only *emm*5 (n=4) and *emm*6 (n=7) were singleton *emm*-cluster types.

**Figure 3:**
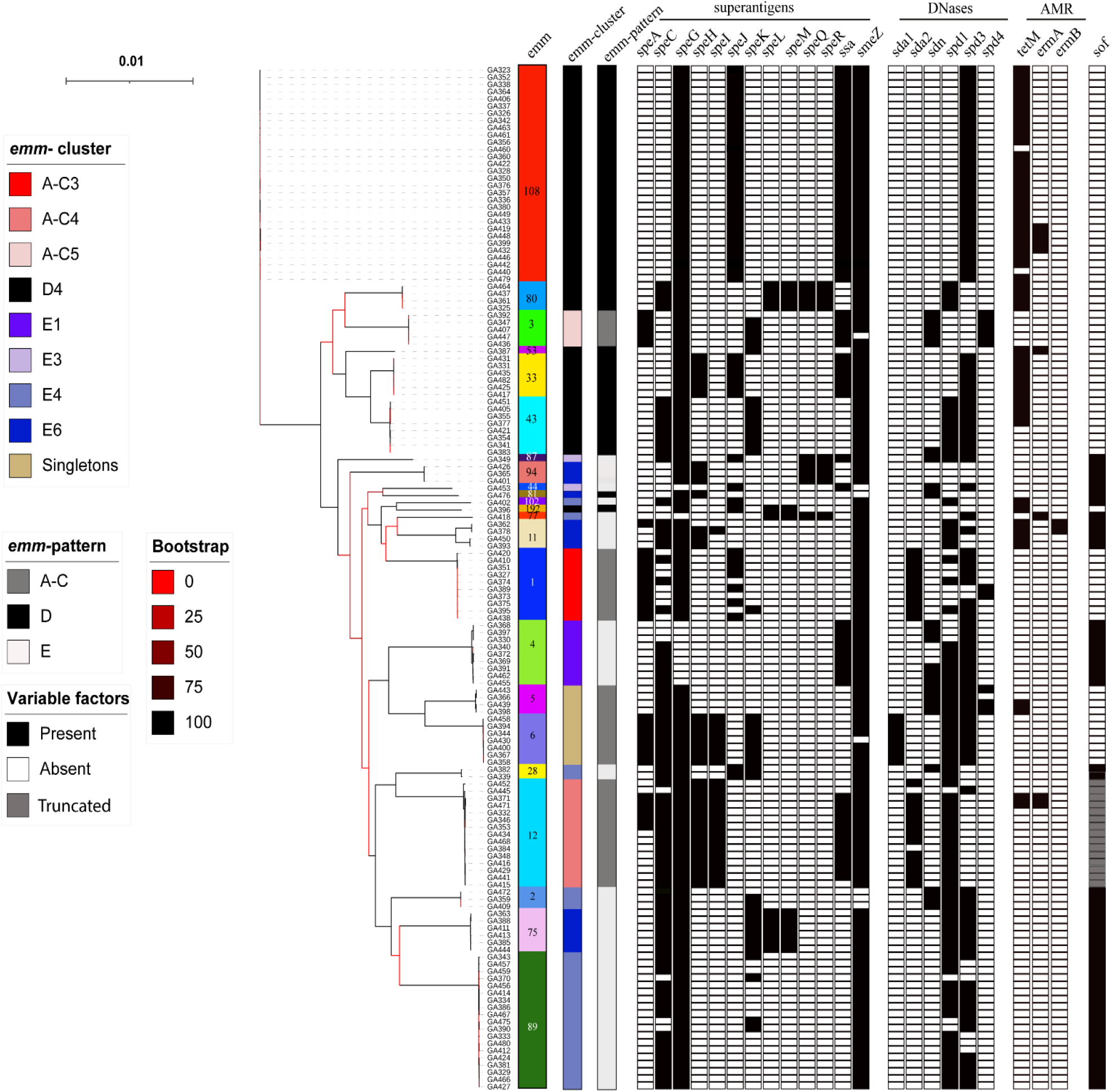
Phylogenetic analysis of 142 HIC isolates: A maximum likelihood phylogenetic tree was generated with the core-gene alignment (1,202,105bp) using RAxML (22) with 100 bootstraps. All isolates clustered by *emm*-type. Presence (black)/absence (white) of superantigens (*speA*, *speC*, *speG*, *speH-M*, *speQ*, *speR*, *ssa* and *smeZ*) and DNases (*sda1*, *sda2*, *sdn*, *spd1*, *spd3* and *spd4*) is indicated. Antimicrobial resistance genes (AMR) *tetM, ermA* and *ermB* were also identified in some isolates (white; absent, black; present). The positivity for serum opacity factor (*sof*) is also shown, but in all *emm*12 this gene would produce a truncated variation of SOF (grey). Scale bar represents substitutions per site. *emm*-types are coloured for easy visualisation and type numbers are also given.

Consistent with the fewer *emm*-type within the HIC isolate collection, we identified only 28 different STs, the most common being ST14, ST101, ST36 and ST28, reflective of their association with the dominant genotypes *emm*108, *emm*89, *emm*12 and *emm*1, respectively. As with the LIC isolates, STs were unique to a single *emm*-type.

The phylogenetic analysis of the HIC isolates based on core-genome SNPs also grouped isolates into lineages based on *emm*-types, and all *emm*-types formed single lineages (Figure 3). Pairwise genetic distance between isolates identified a median of 17 SNPs between isolates of the same *emm* type (range 0 to 2206), compared to a median of 11100 SNPs distance between isolates belonging to different *emm* types (range 3057 to 12339) (Supplementary Figure 1B).

Surprisingly, only seven out of the 62 total *emm*-types identified were common to both LIC and HIC isolates: *emm*4, 28, 75, 77, 80, 81 and 89. However, except for *emm*80 (*emm*80.0), the other six overlapping *emm*-types were of different *emm* sub-types between the two sites (Figure 2, Supplementary table 1). All were *emm*-cluster E *emm*-types, except *emm*80 which belongs to *emm*-cluster D4. Pairwise comparison of isolates from the two different sites within each of these *emm*-types revealed a level of genetic distance similar to that observed when isolates of different *emm*-types were compared, indicating that, although they may share an *emm*-type, they do not share a core genome.

It is also possible that closely related isolates may exist within both collections but carry different *emm* genes. Core-gene phylogeny of all isolates from both sites combined showed clear segregation of isolates from different sites, except in one instance where an *emm*192 HIC isolate clustered with two *emm*56 LIC isolates (Supplementary Figure 2).

The core genome of isolates from both sites combined was 1191 genes from a total of 7921 genes. However, while 1416 genes were present in at least one HIC isolate and absent from all LIC isolates, 3418 genes were present in at least one LIC isolate but absent from all HIC isolates. This indicates a greater accessory genome in LIC isolates. The core genome of LIC isolates alone was 1288, similar to HIC isolates at 1242, but there was a total of 6408 genes in LIC isolates compared to 4411 genes in HIC isolates.

### The Mga*-*regulon diversity

The core Mga-regulon includes the *mga* gene and all intervening genes up to and including *scpA* (encoding for the C5a peptidase). Genes within this region encode proteins involved in cell invasion and immune evasion and include those for the M protein, encoded by *emm*, and the M-like proteins Mrp and Enn. The composition of the intervening genes that define the Mga-regulon, as well as the type of M protein and positivity for serum opacity factor (*sof*), relates to the *emm* pattern (A-C, D or E) (10,28). We were able to determine the composition of the Mga-regulon for 36/46 *emm*-types for 85/107 LIC isolates and all 23/23 *emm*-types for 139/142 HIC isolates. Among the LIC isolates, we could not confirm the Mga*-*regulon for all isolates within ten different *emm* types, because it was not contiguous in the *de novo* assemblies, possibly due to sequence quality or repetitive regions. For the HIC isolates, this was the case for only single isolates within *emm* types *emm*1, 12 and 108, and other isolates within these *emm*-types had confirmed Mga-regulons.

Six different Mga-regulon compositions were identified across isolates from both sites (Figure 4) but the vast majority of *emm*-types from both sites were Mga-regulon type I, consisting of *mga*, *mrp*, *emm, enn* and *scpA*. This type was found in 31/36 *emm*-types in LIC isolates and 16/23 *emm*-types in HIC isolates, accounting for 88% (75/85) and 71% (98/139) of the LIC isolates and HIC isolates, respectively. Mga-regulon type II, with the *emm*1 streptococcal inhibitor of complement (*sic*) or *emm*12 SIC related gene (*drs*), was only found in HIC isolates.

Alleles for *mrp* and *enn* were extracted and compared for associations with *emm* and geographical location of the isolate. Ninety-seven *mrp* genes and 92 *enn* genes were extracted from the 107 LIC isolate genomes, resulting in 44 unique *mrp* sub-alleles and 48 unique *enn* sub-alleles. From the 142 HIC isolate genomes, we extracted 101 *mrp* genes and 99 *enn* genes, resulting in 22 unique *mrp* sub-alleles and 21 unique *enn* sub-alleles. For the majority, unique alleles were associated with *emm*-type and geographical location, although phylogenetic analysis did show overall there was limited geographical restriction between closely related alleles (Supplementary Figure 3). There were two main clades for both Mrp and Enn, each with one clade associated with E cluster *emm*-patterns while the other associated with a mix of *emm*-patterns. We did identify some instances of the same *mrp* allele associated with different *emm* types, although, with one exception, this was restricted to the LIC isolates. The *mrp*202 allele was shared by *emm*119 and *emm*162 isolates and *mrp*60 was shared by *emm*85 and *emm*89 isolates. Sub-alleles (same amino acid sequence but different nucleotide sequence) *mrp*193.14 and *mrp*193.15 were found in *emm*116 and *emm*86, respectively. Different sub-alleles of *mrp*195 were found in the LIC *emm*18, *emm*95 and *emm*/stg1750 isolates but also in HIC *emm*53 isolates. A similar pattern was also found with *enn*, with different sub-alleles of *enn*199 found in the LIC *emm*65 and *emm*182 isolates, and sub-alleles of *enn*26 found in the LIC *emm*168 but also HIC *emm*89 isolates.

**Figure 4:**
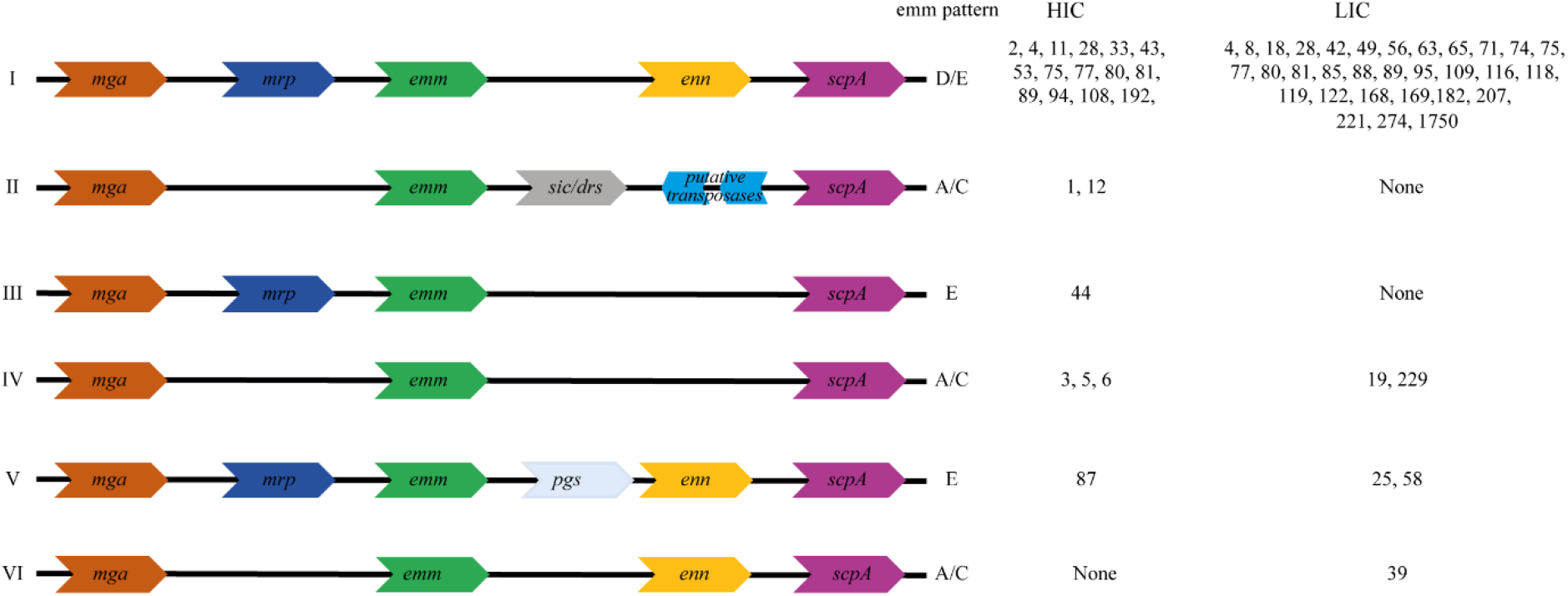
Arrangement of genes in the Mga regulon. The genes within the *mga* regulon for each isolate was determined and an Mga-regulon type I-VI assigned. The majority of *emm* types in both the HIC isolates and LIC isolates had type I with the M-like protein genes *mrp* and *enn* flanking the M protein gene *emm*. The previously assigned *emm* pattern A-C/D/E (based on the *emm* type) is also given. The streptococcal inhibitor of complement (*sic*) gene was only identified in HIC *emm*1 isolates, and the distantly related to *sic* (*drs*) gene found only in HIC *emm*12 isolates. The gene *pgs* encodes for Pgs, a 15.5kDa protein of unknown function (28).

We also looked for the presence of the *fbaA* gene, downstream of *scpA* (outside of the Mga-regulon), which encodes a surface protein associated with the infection potential of pattern D skin isolates (11,31). This gene was found in all D pattern and E pattern isolates but was absent in 75% of A-C pattern HIC and LIC isolates (Supplementary Table 1).

### Diversity of superantigens and DNases in the skin isolates

The complement of superantigen and DNase genes *S. pyogenes* isolates can carry varies, mainly due to the association of these factors with mobile bacteriophages. There are potentially 13 different superantigen genes that can be carried by *S. pyogenes*; *speA*, *speC*, *speH*, *speI*, *speK*, *speL*, *speM*, and *ssa* are prophage-associated, while *speG*, *speJ*, *speQ*, *speR* and *smeZ* are chromosomal. Of the 107 LIC isolates, 99 (93%) carried *speG* and 97 (91%) had *smeZ*. Less common were *speJ* and the co-transcribed *speQ*/*speR*, found in 43/107 (40%) and 7/107 (7%), isolates respectively (Figure 5). A similar pattern was observed in the HIC isolates, with 130/142 (92%) and 134/142 (94%) isolates carrying *speG* and *smeZ* respectively, while *speJ* was present in 46/142 (32%) and *speQ/speR* was carried in 9/142 (6%) isolates.

**Figure 5:**
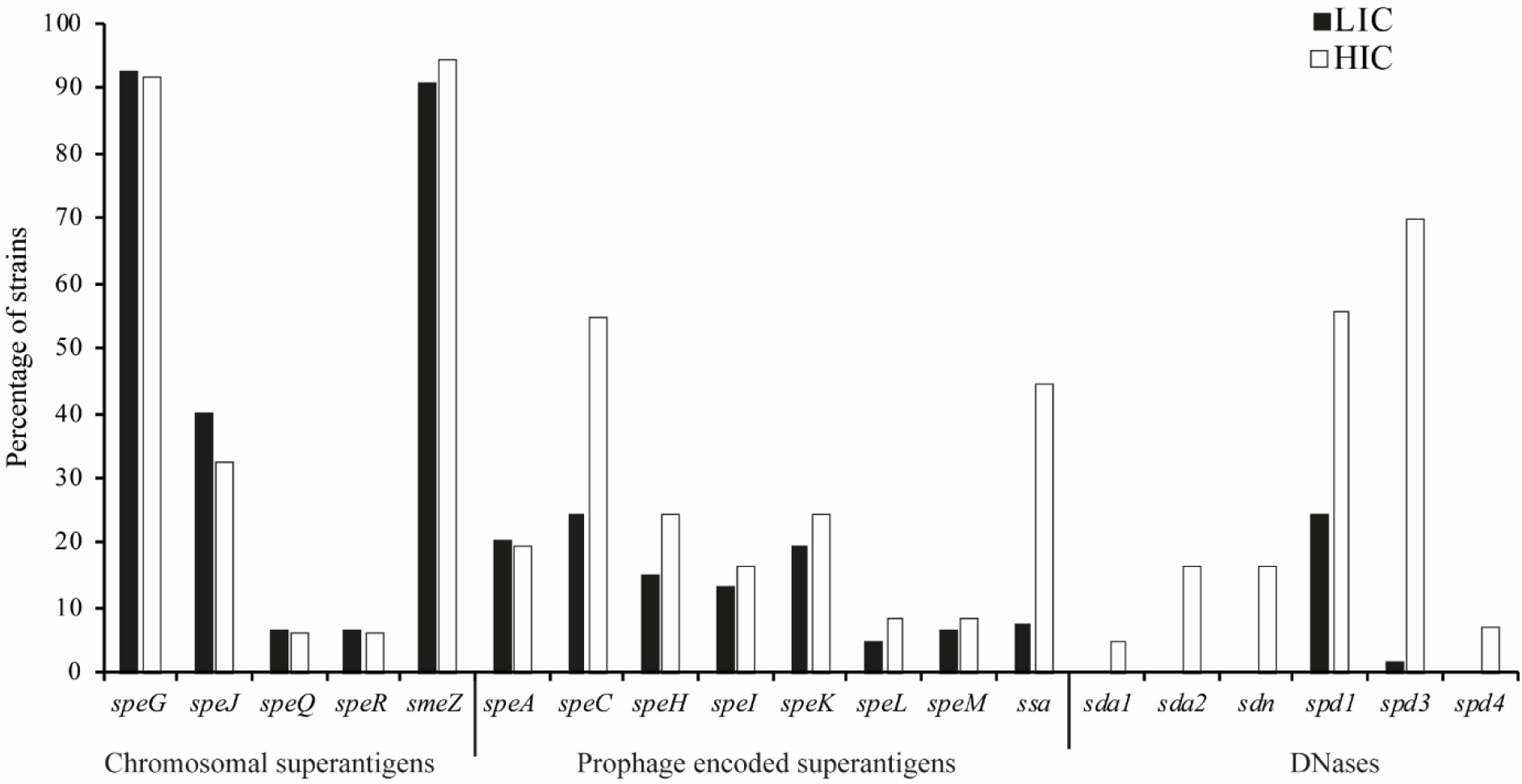
Superantigen and DNase gene carriage in LIC isolates compared to HIC isolates. The proportions of the LIC isolates (black bars) and HIC isolates (white bars) carrying the respective genes determined by BLAST analysis and mapping.

Of the prophage-associated superantigens, *speC* was the most predominant in the LIC isolates, carried by 26/107 (24%) isolates (Figure 5), and in the HIC isolates, although much higher at 55% (78/142). Two (out of eight) *emm*43 HIC isolates and the single *emm*102 HIC isolate each carried two copies of *speC*, as well as two copies of the associated DNase *spd1*. These appeared to be carried on two separate phages integrated at two different sites.

In the HIC isolates, prophage-associated *ssa* was present in 63/142 (44%) isolates, compared to only 8/107 (7%) of the LIC isolates.

Interestingly, *speA* was almost equally common in the LIC isolates (22/107, 21%) as in the HIC isolates (28/142, 20%), but, apart from one *emm*89 isolate, all LIC isolates carried the *speA4* allele (or a *speA* very close to this allele) which is 11% divergent from the other alleles (32) and was associated with a prophage-like element rather than a full prophage. This prophage-like element has been previously identified in the *emm*6 reference strain MGAS10394, termed Φ10394.2, and comprised of transposases and fragments of *speH* and *speI* (Supplementary Figure 4) (32). Previously, it has only been found in *emm*6, *emm*32, *emm*67 and *emm*77. In the HIC isolate collection this element, and the *speA4* allele, was only found in *emm*6. The only isolate in the LIC isolate collection that carried a different *speA* allele, one synonymous base pair different to *speA.1*, was associated with a prophage, although this did not share any substantial identity to other known prophages in *S. pyogenes* (determined by BLASTn against the entire NCBI database).

Prophage-associated *speH*, *speI*, *speK*, *speL* and *speM* were detected at fairly similar levels between the two sites; 15%, 13%, 20%, 5%, and 7% respectively in LIC isolates compared to 25%, 16%, 25%, 8% and 8% in the HIC isolates (Figure 5). One LIC *emm*65 isolate had an apparent fusion gene comprised of 5’ *speK* and 3’ *speM*. An alignment of the 259 amino acids (aa) of this potential fusion protein showed 100% identity to the first 180 aa of SpeK and a 100% of the remaining 181-259 aa to the last 159-237 aa of SpeM (Supplementary Figure 5).

We also tested for the presence of the prophage-associated DNases *sda*, *sdn*, *spd1*, *spd3* and *spd4* (33). Only two prophage-associated DNases were identified in the LIC isolates; *spd1*, 26/107 (24.3%) and *spd3*, 2/107 (1%). These were also the most prevalent in the HIC isolates, at 79/142 (56%) and 99/142 (70%), respectively but we also detected *sda1*; 7/142(5%), *sda2*; 23/147(16%), *sdn* 23/147(16%) and *spd4* 10/142(7%).

### Hyaluronic capsule biosynthesis genes

Although the hyaluronic capsule is considered an important virulence factor, recently it was shown that genotypes *emm*4*, emm*22 and *emm*89 lack the *hasABC* operon required to synthesise the capsule. Additionally, in HICs there is a high proportion of isolates within different genotypes whereby *hasA* or *hasB* has either been deleted or carries a mutation that would render the encoded protein non-functional, predicted to result in the lack or reduction of capsule (33). The *hasABC* operon was detected in all the LIC isolates, including the *emm*4 and *emm*89 isolates, supporting the findings that they have a different core genome compared to HIC *emm*4 and *emm*89, which all lacked the *hasABC* operon. No variations were detected in the *hasA* and *hasB* genes that would lead to truncated proteins in the LIC isolates, except for one *emm*74 isolate with a *hasA* variant that would encode for a truncated HasA. In the HIC isolates and consistent with previous findings (33), all *emm*28, *emm*77 and *emm*87 isolates were predicted to produce truncated HasA, and all *emm*81 and *emm*94 predicted to produce truncated HasB. Three other isolates were predicted to produce truncated HasA and a further two to produce truncated HasB, but these were sporadic examples within *emm*-types (Supplementary Table 1).

### FCT-types in the LIC and HIC isolates

The Fibrinogen collagen binding T-antigen (FCT) region, which is classified into 9 different types (FCT1-9), encodes for pilin structural and biosynthesis proteins and adhesins that could be potential determinants of genetics basis for tissue tropism (34). Therefore, we investigated the diversity of the FCT regions in isolates across the two geographical settings. Eight different patterns were identified across the two sites, corresponding to FCT1-6 and FCT9, as well as a previously unidentified pattern found among the LIC isolates, which we termed FCT10; it was similar to FCT5, but with an additional fibronectin binding protein (Figure 6). FCT3 was found in the most *emm*-types in both LIC and HIC isolate collections, 9/23 (39%) and 20/46 (43%), respectively, although this represented only 23% of the HIC isolates compared to 41% of LIC isolates. FCT4 was also found in a high proportion of *emm* types, accounting for 7/23 (30%) and 11/46 (24%) *emm*-types, representing 28% and 30% of HIC and LIC isolates, respectively. Due to the prevalence of *emm*108 and *emm*1 in HIC, 33% of isolates were either FCT1 or FCT2, whereas only 6% of the LIC isolates were FCT1 and no LIC isolates were FCT2. There was only one example of isolates of the same *emm*-type with two different FCT-types, and that was within the two LIC *emm*118 isolates. While one *emm*118 (ST1205) isolate was estimated to be FCT4, the other (ST354) was estimated to be FCT10, alongside the two LIC *emm*63 isolates. The FCT regions in both *emm*118 isolates however were estimated as they were not found within a single contiguous sequence.

**Figure 6:**
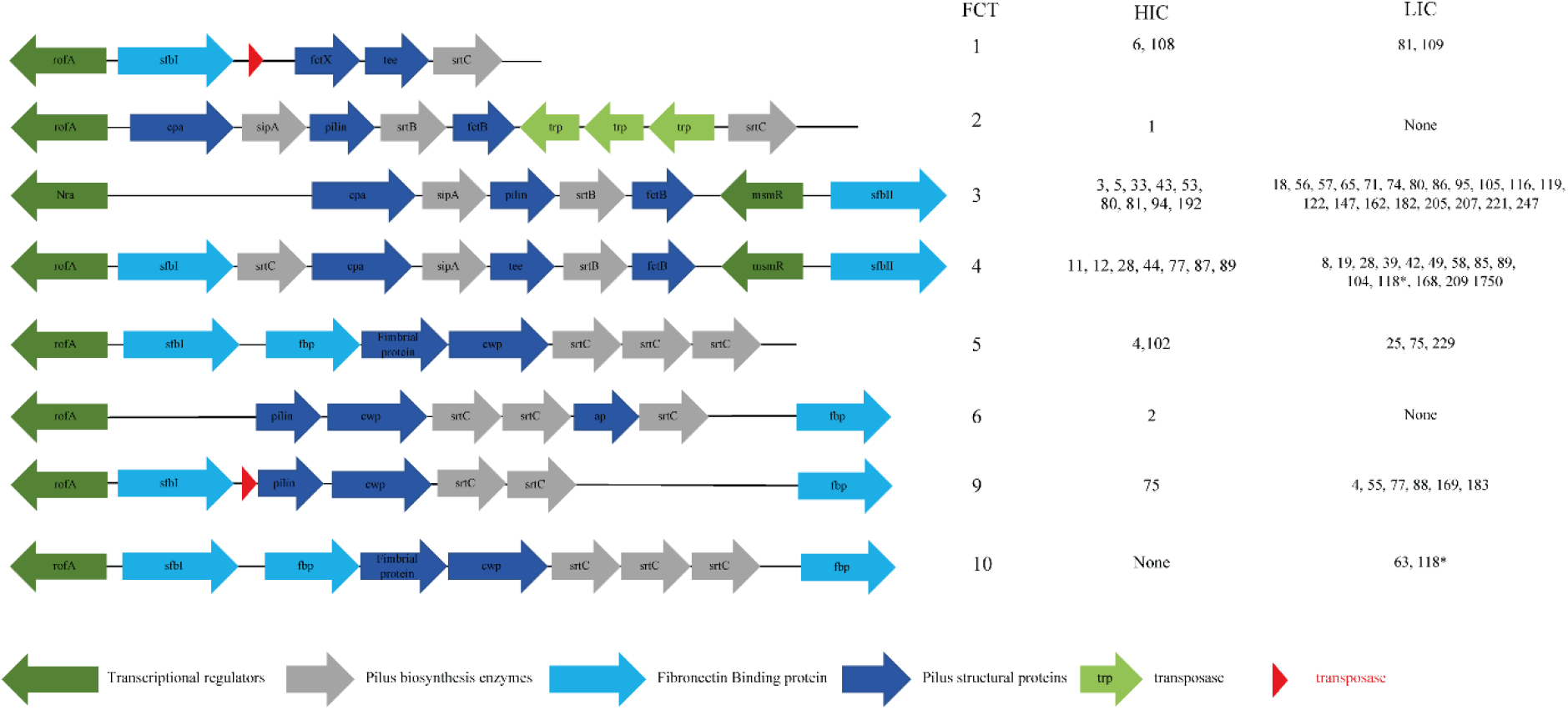
FCT arrangement patterns identified in LIC and HIC *S. pyogenes* isolates. FCT regions were extracted from *de novo* assemblies and the FCT type assigned based on the predicted function and order of genes within the extracted region. The *emm*-types of isolates with each FCT type are shown for HIC and LIC isolates. A new FCT region was identified (FCT10) as similar to FCT5 but with an additional fibronectin binding protein after the sortase genes. For all *emm*-types there was at least one isolate with a designated FCT type in a single contiguous region. The only exception to this was *emm*118 (*) where the FCT was estimated to be FCT4 and the new FCT10 for each of the two isolates as the FCT region was split over two contigs. In FCT1 transposases were found in HIC *emm*6 and *emm*108, and in FCT9, transposases were found in HIC *emm*75 and LIC *emm*4. fbp; fibronectin binding protein, cwp; cell wall protein, ap; ancillary protein and trp; transposase.

We also compared the amino acid sequences of the FCT regulatory genes *rofA*, *nra* and *msmR* and identified a number of different of variations. For the majority, variations were common to all isolates within an *emm*-type and there were no obvious variations that may affect function. We found that 9/10 HIC *emm*1 isolates carried three variations within RofA that characterised them as being part of the M1UK lineage associated with high *speA* expression (35). No other isolates were found to carry any of these three RofA variations.

### Prevalence of antimicrobial resistance genes

Of the 107 Gambian assemblies, the *tetM* gene encoding for tetracycline resistance was identified in 79/107 (73.8%), and 37 of these (33.6% of the total population) also carried the *tetL* gene and one carried *tetK*. Furthermore, *dfrG* or *dfrK*, both encoding for trimethoprim resistance, were identified in 10/107 (9.3%) and 17/107 (15.9%) of isolates respectively. Only 53/142 (37.3%) of the HIC isolates carried the *tetM* gene (Figure 1 and 3) and no other resistance genes were found except for *ermA* in 8/142 (6.5%) isolates and two *emm*11 isolates carried *ermB, sat4A and aph3*.

### Vaccine antigen diversity

Based on the number of isolates with *emm*-types present in the vaccine, the potential coverage of the 30-valent M protein vaccine in the LIC isolates was 24%, with only 11 vaccine-included *emm*-types (Supplementary Figure 6). On the other hand, the potential coverage of the HIC isolates was 61%, although only 14 were vaccine-included *emm*-types. This suggests limited potential for this vaccine for low-income settings such as The Gambia, although there may be potential for cross-protection as has been seen for some *emm*-types (4, 36).

Among other potential vaccine candidates, the genes *spy0651*, *spy0762*, *spy0942*, *pulA*, *oppA*, *shr*, *speB*, *adi*, *ropA*(*tf)*, *spyCEP*, *slo*, *spyAD*, *fbp54* and *scpA* were recently highlighted as conserved potential targets (37). All LIC and HIC isolates carried all 14 genes and BLASTp indicated that all genes were highly conserved in all isolates with less than 1% sequence divergence (>99% identity) from the corresponding genes in reference genome MGAS5005 (*emm*1).

## Discussion

The overall global burden of *S. pyogenes* infection and associated post-infection sequalae, highlights the need for more research into treatment and prevention, with a particular focus on vaccine development. Maximal global impact of a preventative vaccine against *S. pyogenes* can only be achieved on the back of better understanding of the global diversity of the *S. pyogenes* population, but to date, large-scale genomic studies have been mainly focused on HIC isolates. The Gambia, West Africa is a LIC with a high burden of streptococcal skin infections (15). Studies on circulating *emm*-types in this region, and in other African countries, indicate a much higher level of diversity than that seen in HICs (6–9) and this is reflected in the limited African genomic data (37). In this study, we aimed to contribute genomic data and provide molecular characterisation of *S. pyogenes* in The Gambia by whole genome sequencing isolates collected during a population-based study of skin infections in children aged 5 years and under. To act as a comparison isolate collection, we also genome sequenced isolates from Sheffield, UK to represent HIC isolates.

Consistent with other findings from LICs (9), we identified a high number of different *emm*-types in the LIC isolate collection from The Gambia compared to the HIC isolate collection from the UK, and no dominant type. In the HIC isolates, five *emm*-types (*emm*108, *emm*89, *emm*12, *emm*1 and *emm*4) accounted for ~60% of the isolates. There was also limited overlap across the two sites with only 7 shared *emm*-types; *emm*4, 28, 75, 77, 80, 81 and 89. However, it was clear that these *emm-types* represented a different genetic background between the two locations, supporting previous findings that *emm* might not be a good marker for characterising a diverse global population (37).

Although we did not specifically select for impetigo isolates or patient age range amongst the HIC isolate collection, all were associated with some form of non-invasive skin infection. Little molecular information is available for *S. pyogenes* causing skin infections in the UK, as isolates are not routinely collected and typed, or for other HICs. The dominant *emm* genotypes found in the HIC isolates reflected what has been found in other types of infections, with *emm*1, *emm*12 and *emm*89 leading among invasive isolates in the UK (33) and *emm*1, *emm*4, *emm*12 and *emm*89 common among UK scarlet fever cases and upper respiratory tract infections (35, 38). Very similar patterns of *emm*-types causing invasive disease are also found in other European countries and North America, with *emm*1, *emm*28, *emm*89, *emm*3, *emm*12, *emm*4 and *emm*6 leading (39). The genotype *emm*108 has not previously been reported to be a common *emm*-type in the UK or elsewhere, but reported in 2018/2019 by Public Health England to be a cause of national upsurges in infections in England/Wales (https://assets.publishing.service.gov.uk/government/uploads/system/uploads/attachment_data/file/800932/hpr1619_gas-sf3.pdf). The data on prevalence of this *emm*-type is based on invasive disease data, as only invasive infections are notifiable in England/Wales. From the available data it is not clear if it would have been common among throat infections during this time as well as skin infections, but suggests it is not unique to our sampled geographical region of Sheffield, UK.

The *emm*-pattern D, previously determined to be associated with skin infections, was the most common in the LIC isolates (48%) and the HIC isolates (36%), although *emm*-pattern E was almost equally as common in HIC isolates (35%). A review of population-based studies (11) found that among impetigo isolates, 49.8% were D, 42% were E and 8.2% were A-C patterns, compared to 1.7% D, 51.7% E and 46.6% A-C patterns among pharyngeal isolates. This distribution is consistent with our findings in the LIC isolates (48% D, 40% E, 12% A-C) but we found a higher level of A-C isolates (29%) in HIC isolates. This could be due to the more diverse collection of HIC isolates, given that we did not focus specifically on impetigo. Interestingly, the dominant HIC *emm*-types were either pharyngeal specialist pattern A-C (*emm*1 and *emm*12) or generalist pattern E (*emm*4 and *emm*89), with only *emm*108 representing skin specialist pattern D.

In the LIC isolates, all six E *emm*-clusters were represented, with the most common being E6 (18%) closely followed by E4 (16%) and E3 (14%). E6 was recently found to be the leading cluster in Gambian non-invasive isolates (skin and pharyngeal) but with E3 leading among invasive isolates (9). D4 was also common in LIC isolates (17%) but, more so in HIC isolates where 35% of the isolates were D4. This was almost equal to all E clusters combined, but again explained by the high number of *emm*108 isolates. A higher number of singleton *emm*-cluster types were also found in the LIC isolates (n=17) representing 9 *emm*-types, compared to HIC isolates (n=11) representing just two *emm*-types. There was an association with E *emm*-cluster isolates also carrying the *sof* gene, as all E1-E4 *emm*-types were *sof* positive. Four LIC E6 *emm*-types (*emm*46, 65, 182 and 205) were *sof* negative and all E5 *emm*-types were negative. HIC *emm*12 isolates carried a *sof* gene that would only produce a truncated form of SOF, as previously identified (11).

Consistent with the high number of D/E pattern isolates, we also found the majority of isolates had the Mga-regulon pattern I, and therefore carried the *emm*-like genes *mrp* and *enn*. Within the HIC *emm*4 isolates we found that 4/9 carried the *emm*-*enn* fusion gene, and this was also associated with degraded prophages in these isolates (40, 41). Given the high number of isolates carrying Mrp and Enn it is possible that they contribute to pathogenesis at the same, or even greater, level of the M protein (28). The M-like proteins have not been well characterised and their role and expression may vary depending on the allele or other genetic factors. The existence of two major clades within the Mrp and Enn phylogeny is of interest and may indicate varying domains and functions. Despite being adjacent to the *emm* gene, we did not observe sharing of *enn* and *mrp* alleles with *emm*-type over the two geographical sites. We did, however, see the same allele or very closely related alleles of *mrp* and *enn* shared with different *emm*-types across different geographical locations.

HIC *emm*4 and *emm*89 isolates were acapsular, as expected, but this was not the case for LIC *emm*4 and *emm*89, again reflecting very different genetic backgrounds. All LIC isolates carried the *hasABC* genes required to synthesise the capsule, only one isolate had a mutation that would lead to a truncated HasA and a probably acapsular phenotype.

The FCT region encodes for genes thought to be involved in adhesion to the host, particularly the pili, which are likely to mediate primary host:pathogen interactions (42). Factors essential for pili construction are encoded within the FCT and include a major pilus subunit, one or two minor subunits, at least one specific sortase and a chaperone (42). The pili of the M1 isolate, SF370, has been shown to be essential for adherence to human tonsil and human skin (43), indicating its role in primary interactions and establishing infection. Other factors included within the FCT region are fibrinogen and fibronectin binding proteins, which may also contribute to host cell interactions, as well as transcriptional regulators. We identified the previously described FCT types FCT1-6 and FCT9 among our isolates but, also a new FCT type (FCT10) that was based on FCT5 with an additional fibronectin binding protein. FCT2 and FCT6 was restricted to HIC isolates and the new FCT10 was only found in LIC isolates.

FCT3 and FCT4 were the most common types across both sites, found in 70% (16/23) and 74% (34/46) of *emm*-types, representing 54% (76/142) and 69% (74/107) HIC and LIC isolates, respectively. FCT3 and FCT4 have been shown to share the greatest similarity and can undergo recombination (42). Both these FCTs have a *cpa* gene, which encodes for a collagen binding subunit found at the pilus tip, one or two fibronectin-binding proteins (*sfbI/sfbII*) and the regulator *msmR* upstream of the fibronectin-binding protein. The pilus and fibronectin-binding proteins may contribute to tissue-specific host cell adhesion, in addition to others located outside the FCT region. This includes *fbaA*, which we identified to present in all isolates except for the majority of A-C pattern types, and has been found to contribute to skin infection (31). The regulator *msmR* has been shown to have a positive effect on the fibronectin binding protein expression and may also control other surface proteins, impacting on host cell adhesion (44). It is not clear if specific FCT types confer tissue tropism and previous work has shown that there is a high level of variability in host cell interactions and biofilm formation between isolates sharing the same FCT (45). This indicates that there are other bacterial factors involved in the expression of FCT related genes. The role of the regulators *nra* or *rofA* do vary between isolates of differing genetic backgrounds, with evidence of environmental effects such as pH and temperature (42). We explored the sequences of *rofA, nra* and *msmR* and found a number of different variations, however, many seemed to be related to *emm*-type and it is difficult to determine if any variation would impact on function. This was also the case for the two-component regulator CovR/S and the regulator of *cov*, RocA, for which variations can impact on the expression of a number of virulence factors. Variations in CovS and RocA were common among both LIC and HIC isolates but the transcriptional impact of any of these amino acid changes is unclear. Only one HIC isolate had an amino acid difference in CovR (M17I, *emm*77) and one other HIC isolate had a premature stop codon in CovS; both may alter expression of virulence genes. Whether there are differences in expression and control of FCT and other virulence factor genes in LIC isolates compared to HIC isolates and/or between skin infection isolates and other types of infection isolates is yet to be determined. Inclusion of isolates causing other infections, such as pharyngeal infection isolates may reveal some tissue tropism differences or factors. However, the complex nature of regulatory systems also makes it difficult to determine the impact of single amino acid variants and control of transcription may vary between *emm*-types.

Superantigens are important *S. pyogenes* virulence factors and their distribution may differ between isolates. The chromosomal *speG* and *smeZ* genes were the most common in both populations, with more than 90% of the isolates carrying these genes. The prophage-associated *speC* and *ssa* were more common in HIC isolates compared to LIC isolates, and three HIC isolates actually carried two copies of *speC*, along with the DNase *spd1*, on two separate prophages. Typically, *speA* is prophage associated but the divergent *speA*.*4* allele is associated with a prophage-like element that has been previously only found in in *emm*6, *emm*32, *emm*67 and *emm*77 (32). We found this only in the HIC *emm*6, but, although *speA* was almost equally as common in the LIC population, all, except one, of the 22 *speA*-positive LIC isolates carried *speA*.*4* associated with the prophage-like element. Only a LIC *emm*89 isolate carried *speA* on what appeared to be a complete prophage and was only one base pair different from the *speA*.*1* allele. Interestingly, we also identified a gene in one LIC isolate (*emm*65) that appeared to be a fusion of 5’ *speK* and 3’ *speM*, and since *speK* and *speM* are phage encoded, it could be a result of recombination of phages carrying the two genes. BLASTp of this potential fusion protein identified a similar (two-three amino acid different) variant in six published genomes; NS88.3 (*emm*98, locus accession PWO34032), *emm*89.14 (QCK42181), *emm*100 (QCK70992), NS426 (VGQ95836), NS76 (VGR28970) and NS6221 (VHG25078).

Only two of the prophage-associated DNases (*spd1* and *spd3*) were found in the LIC isolates, while five DNases (*sda1*, *sda2*, *sdn*, *spd1*, *spd3* and *spd4*) were identified in the HIC isolates. Almost all (136/142, 96%) of the HIC population carried at least one prophage-associated DNase, whereas only two LIC isolates carried *spd3* and only 24% of isolates carried *spd1*, which associated with the superantigen *speC*. DNases, such as *sda1*, have been shown to be necessary and sufficient to degrade neutrophil extracellular traps (46), therefore the lack of these in LIC isolates from The Gambia could be suggestive of limited/reduced ability of immune evasion, and warrants further investigation into their invasive capacity. There is the potential that other prophage-associated DNases exist but are yet to be identified. It also suggests differences in circulating phages between the two sites, although the accessory genome appeared to be much greater in LIC isolates compared to HIC isolates. This could be related to the high prevalence of tetracycline resistance genes within the LIC population that may be carried on mobile genetic elements. Further investigation is needed to determine prophage content, as well as other mobile genetic elements; this is, however, notoriously difficult with short read sequence data and may require supporting long read data.

The most advanced multi-valent *S. pyogenes* experimental vaccine is based on 30 *emm*-types identified from isolates causing infection predominantly in high income countries (4, 5). Based on the *emm*-types distributions, we determine the direct coverage of the vaccine to be only 24% in the LIC population, compared to 61% in the HIC population, although we did not explore cross-reactivity between *emm*-types. The high proportion of *emm*108 in HIC isolates was unexpected as this was not a previously recognised dominant *emm*-type and highlights the potential for sudden and dramatic increases in new *emm*-types that could escape a serotype-specific vaccine. If such a vaccine was introduced, monitoring of new variants in the non-invasive as well as the invasive bacterial populations would be needed, and on a global scale. Alternatively, a vaccine targeting antigens with limited variability between isolates may be preferable, if these can still provide similar levels of protection. We have confirmed that several previously identified potential targets (37) are also highly conserved in our LIC and HIC bacterial populations. However, both our LIC and HIC isolates represent only single geographical locations: Sukuta, The Gambia and Sheffield, UK. Further in-depth genomic analysis of international *S. pyogenes* populations, encompassing more LICs and different infection types, is needed to confirm diversity and distribution of potential vaccine diversity.

Our study confirms work by others (37), that *emm*-typing alone is insufficient to comprehensively characterise global isolates. Furthermore, genetic features that have been characterised in particular HIC *emm*-types, such as the absence of the *hasABC* locus in *emm*4, may not be present in LIC isolates of the same genotype. In the absence of WGS, other molecular markers, such as MLST, *enn*, *mrp* and FCT type could be used in addition to *emm*-typing to characterise the diverse genetic background of isolates from different geographical settings. More work is required to understand why there is such a high genetic diversity in LIC settings compared to HIC and with limited overlap. This may be linked to infection types but there is insufficient data both on pharyngeal infections in LICs, like The Gambia, as well as skin infections in HICs. By increasing the characterisation of isolates from different infections over wider geographical settings we could gain real insight into the molecular mechanisms underpinning tissue tropism.

## Contributors

E.P.A, M.M and T.I.d.S coordinated collection of the Gambian isolates; L.T coordinated collection of the UK isolates; S.Y.B, A.J.K, E.S, S.D, L.T and H.K cultured bacterial isolates and extracted genomic DNA; J.M and A.K.S performed the whole genome sequencing of the Gambian isolates; S.Y.B and C.E.T performed the whole genome sequencing analyses with assistance from R.R.C.; S.Y.B, C.E.T and T.I.d.S secured funding for the project; S.Y.B and C.E.T wrote the manuscript. All authors reviewed and edited the manuscript.

## Acknowledgements

We would like to thank the invaluable contribution of the members of the MRCG Strep A Study Group whose names are not in the main authorship list: Annette Erhart; Pierre R Smeesters; Martin Antonio; Sona Jabang; Beate Kampmann; Anna Roca; Isatou Jagne Cox; Peggy-Estelle Tiencheu; Grant Mackenzie.

This work is supported by Global Challenge Research Fund obtained through the University of Sheffield (S.Y.B). The authors also thank the MRCG at LSHTM and study participants. C.E.T is a Royal Society & Wellcome Trust Sir Henry Dale Fellow (208765/Z/17/Z). T.I.d.S is supported by a Wellcome Trust Intermediate Clinical Fellowship (110058/Z/15/Z). E.P.A is supported by a Wellcome Trust Clinical PhD fellowship in Global Health (222927/Z/21/Z). The authors thank Prof Pierre Smeesters and Dr Anne Botteaux (Universite Libre de Bruxelles) for kindly providing us with new allele numbers for *enn* and *mrp*.

## Supplementary Figures

**Supplementary Figure 1:**
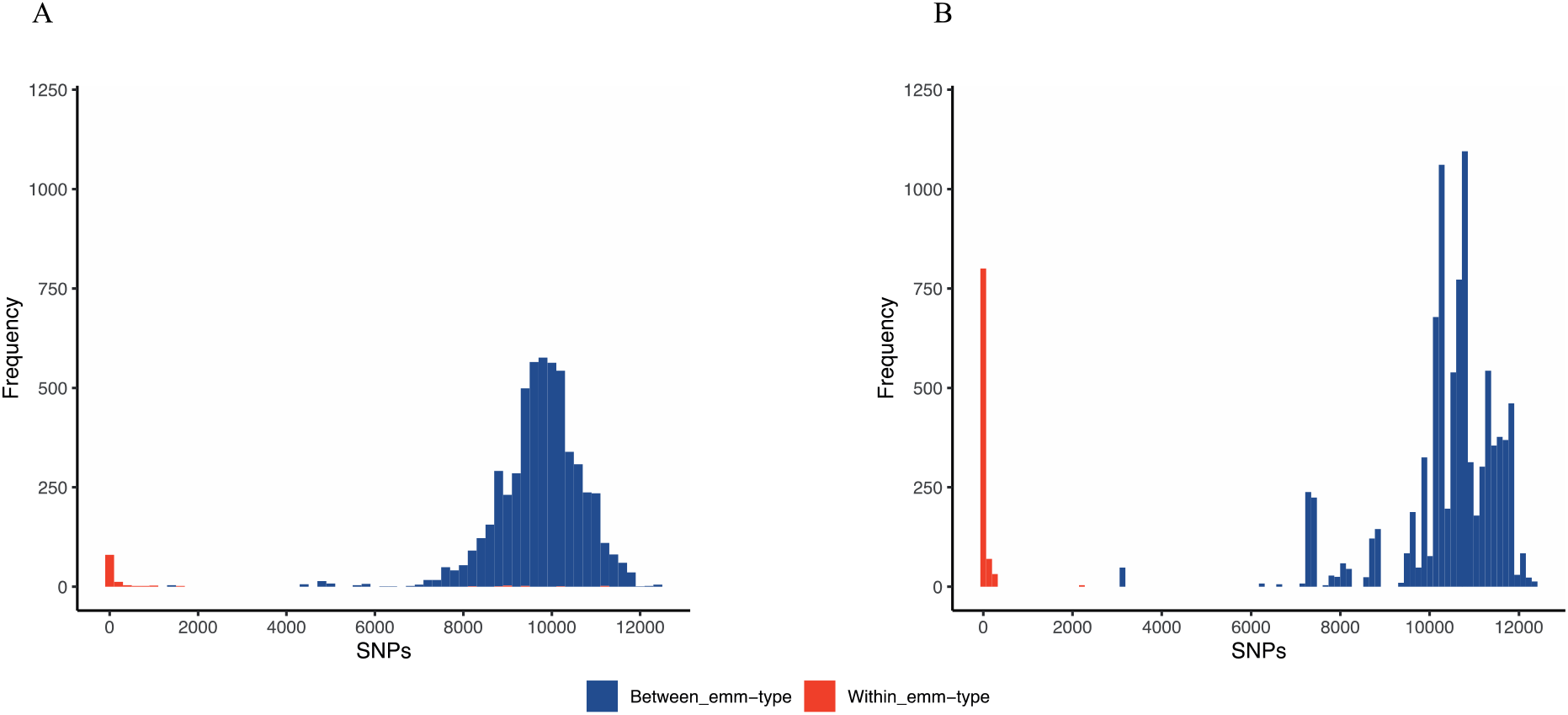
Pairwise single nucleotide polymorphisms (SNPs) distances. SNPs were determined from the core-genome of (**A**) 107 LIC isolates and (**B**) 142 HIC isolates and pairwise distance calculated between isolates belonging to the same (red) or different (blue) *emm*-type. Overall, the median pairwise SNP distance within the same *emm*-type of LIC isolates was 22 (range 0-11,142 SNPs), similar to that of HIC isolates with a median of 17 (range 0-2,206). Also comparable was the between *emm*-type median SNP distance; 9,816 (range 1,423-12,428) for LIC isolates, 11,110 (range 3,057-12,339) for LIC isolates.

**Supplementary Figure 2:**
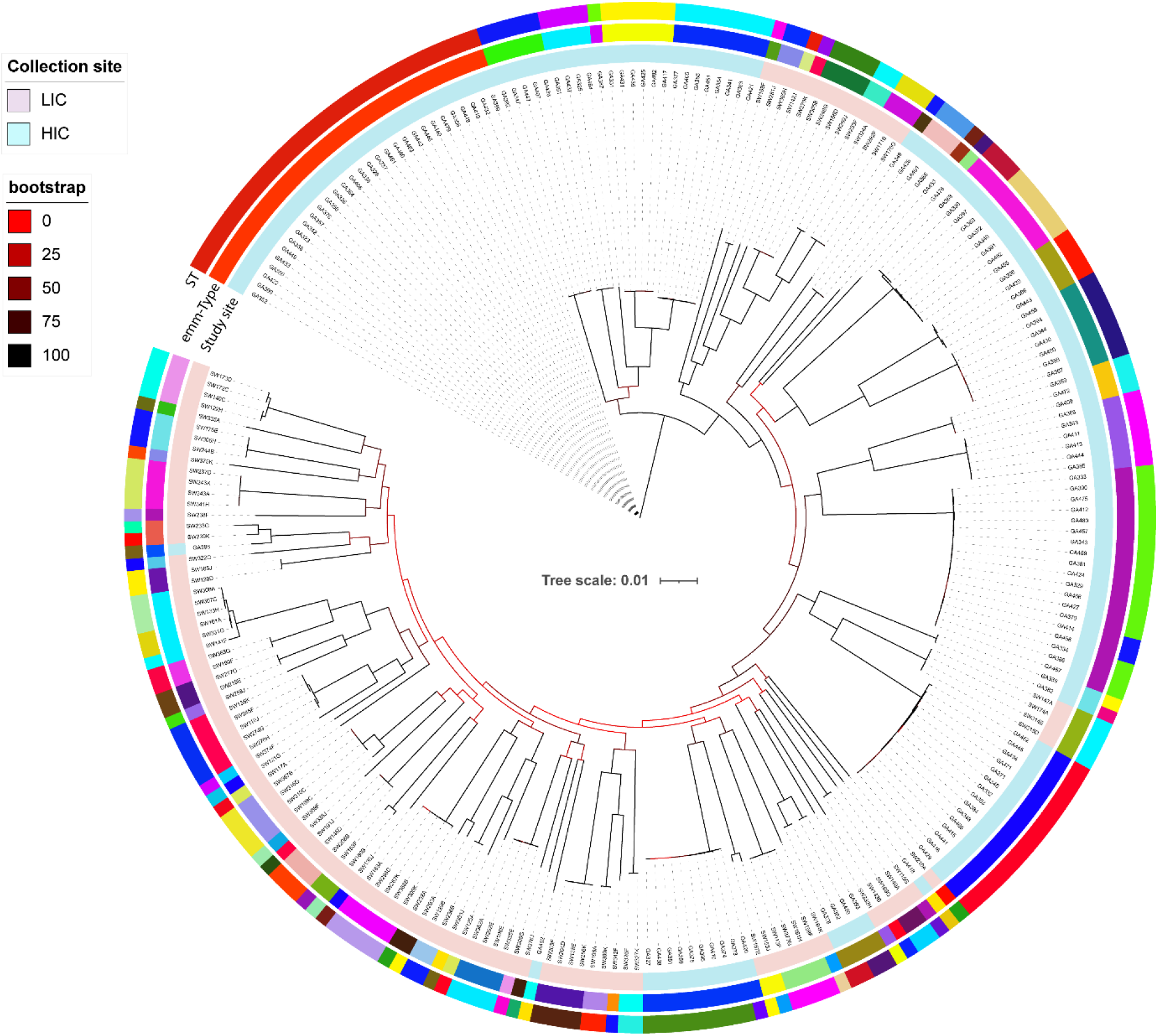
Population structure of the combined LIC (107) and HIC (142) isolates. A maximum likelihood phylogenetic was generated from the core-gene alignment (1,146,086bp) using RAxML with 100 bootstraps. Bootstrap support is indicated by colours in the legend. Inner circle: site of collection, middle circle: *emm*-types and outer circle: ST.

**Supplementary Figure 3:**
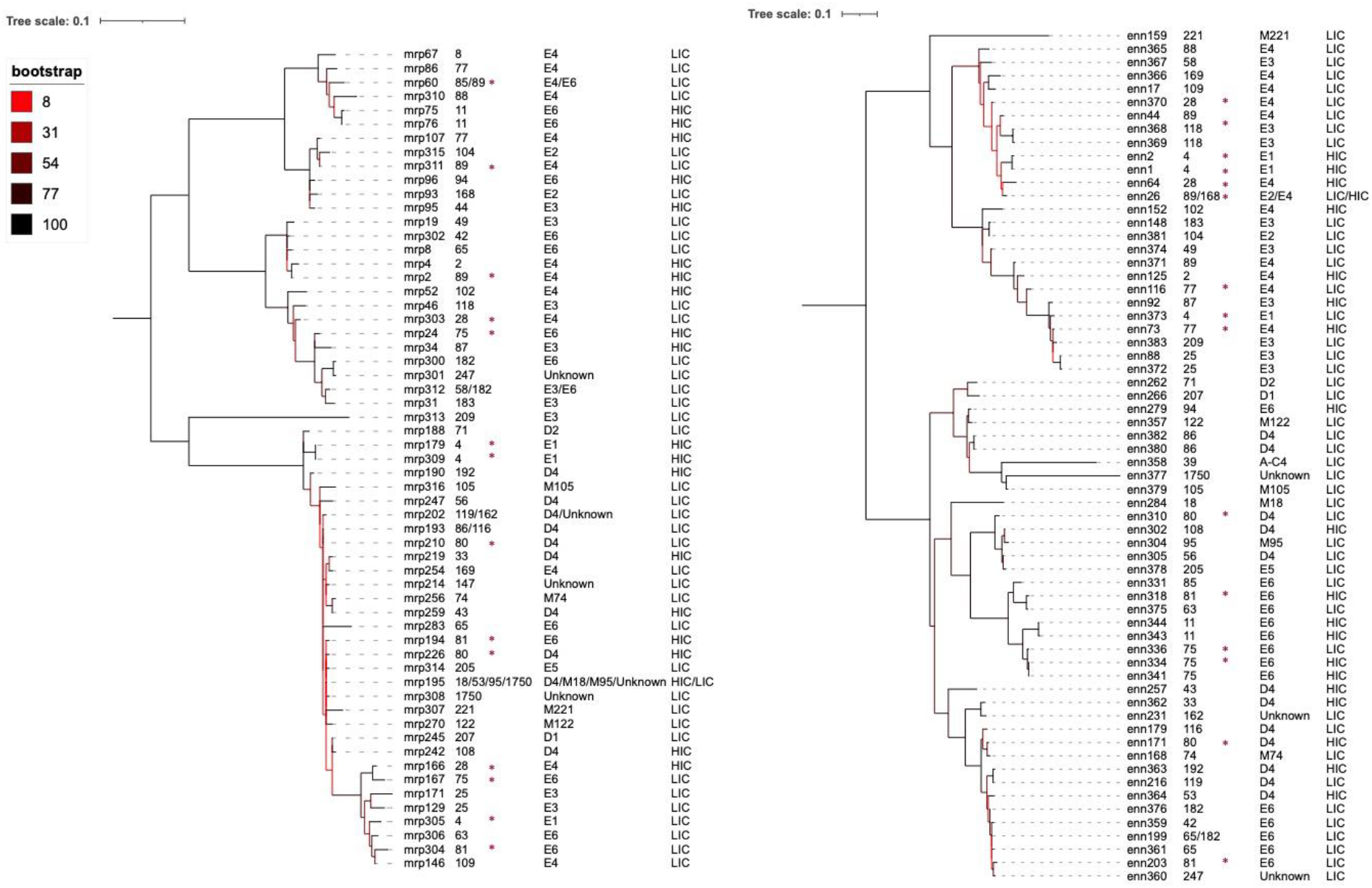
Phylogenetic relatedness of unique Mrp (A) Enn (B) alleles. A maximum likelihood phylogenetic tree was generated from an amino acid alignment of unique Mrp or Enn alleles, using RAxML with 100 bootstraps (branch support shown by colour scale). The Mrp or Enn allele is shown followed by the associated *emm*-type(s), *emm* cluster(s) and population (LIC or HIC). * indicates the shared *emm*-types identified in both sites.

**Supplementary Figure 4:**
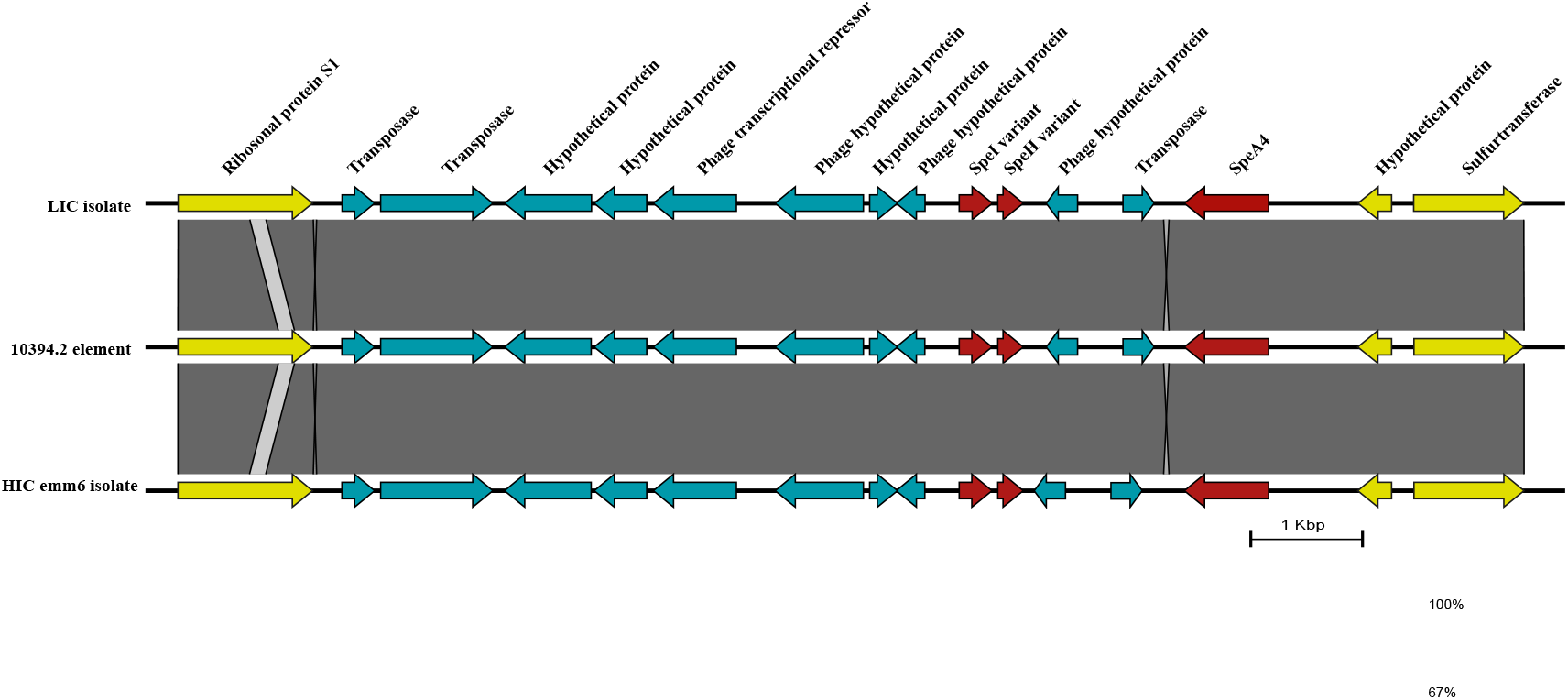
Comparison of 10394.2 phage-like element with the region found in HIC *emm*6 isolates and LIC isolates. The *speA4* in LIC isolates and HIC *emm*6 were located within this phage-like element. This region also contains fragments of *speI* and *speH*. The same element was found in all LIC isolates that carried *speA*, except one that carried a different *speA* allele associated with a prophage. The corresponding regions were extracted from the respective isolate genomes and figure generated using EasyFig.

**Supplementary Figure 5:**
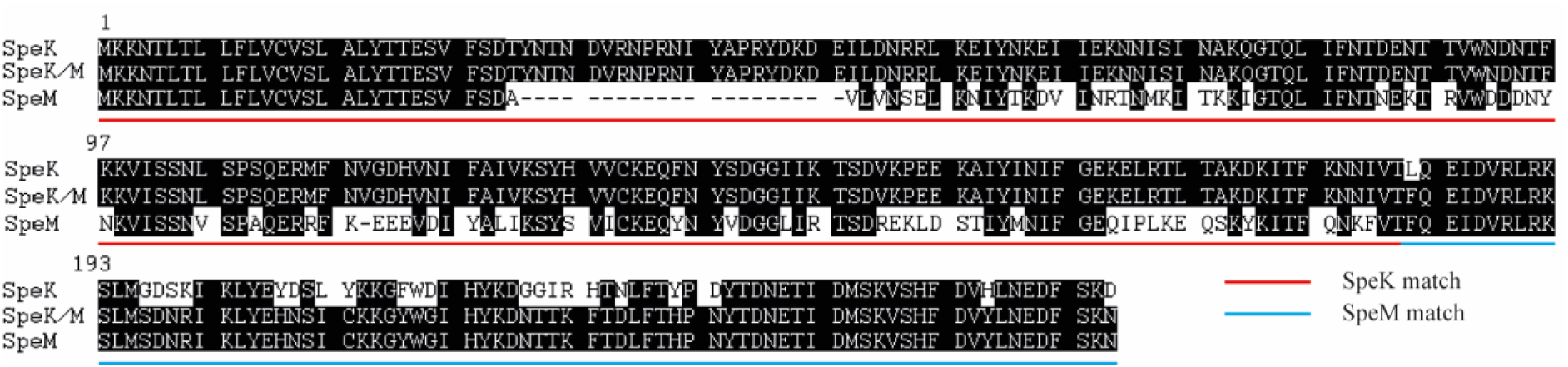
Alignment of the SpeK/SpeM fusion protein to SpeK and SpeM. Within an *emm*65 isolate from LIC, we identified a gene that encodes for 259 amino acids (aa) of which the first 180 aa were 100% identical to the first 180 aa of SpeK (red underlined) but the remaining 181-259 aa were 100% identical to the last 159-237 aa of SpeM (blue underlined). Black shading indicates identical aa.

**Supplementary Figure 6:**
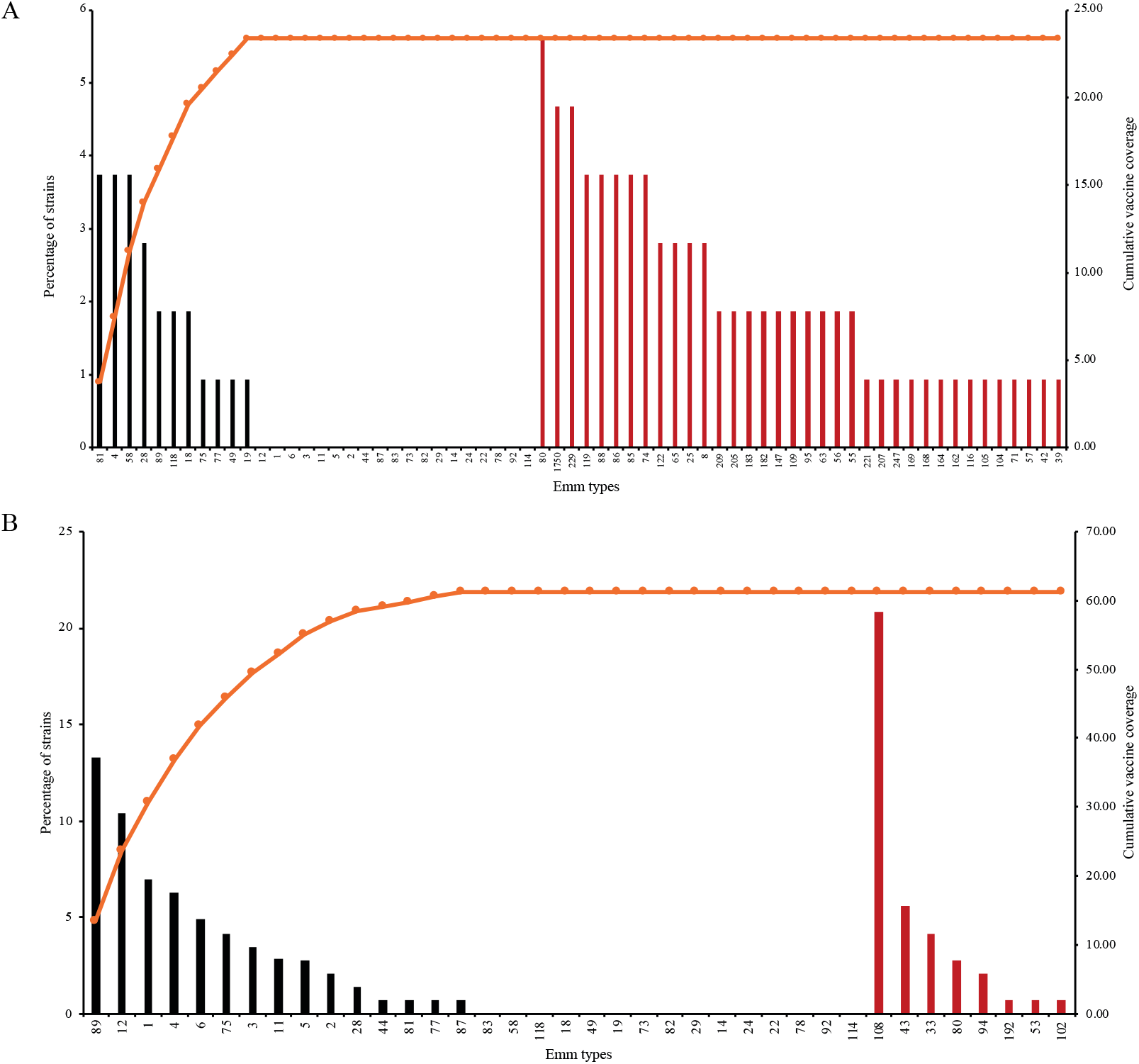
Potential coverage of the *S. pyogenes* 30-valent vaccine. The percentage of (A) LIC and (B) HIC isolates of *emm*-types included in the 30-valent vaccine (black) and other *emm*-types identified in each site but not included in the vaccine (red). The cumulative vaccine coverage for each site is also shown. The *emm*-types without the bars are vaccine included *emm*-types but not seen in the dataset.

